# Unsupervised Neural Network Models of the Ventral Visual Stream

**DOI:** 10.1101/2020.06.16.155556

**Authors:** Chengxu Zhuang, Siming Yan, Aran Nayebi, Martin Schrimpf, Michael C. Frank, James J. DiCarlo, Daniel L. K. Yamins

## Abstract

Deep neural networks currently provide the best quantitative models of the response patterns of neurons throughout the primate ventral visual stream. However, such networks have remained implausible as a model of the development of the ventral stream, in part because they are trained with supervised methods requiring many more labels than are accessible to infants during development. Here, we report that recent rapid progress in unsupervised learning has largely closed this gap. We find that neural network models learned with deep unsupervised contrastive embedding methods achieve neural prediction accuracy in multiple ventral visual cortical areas that equals or exceeds that of models derived using today’s best supervised methods, and that the mapping of these neural network models’ hidden layers is neuroanatomically consistent across the ventral stream. Moreover, we find that these methods produce brain-like representations even when trained on noisy and limited data measured from real children’s developmental experience. We also find that semi-supervised deep contrastive embeddings can leverage small numbers of labelled examples to produce representations with substantially improved error-pattern consistency to human behavior. Taken together, these results suggest that deep contrastive embedding objectives may be a biologically-plausible computational theory of primate visual development.

---

The remarkable power of primate visual object recognition is supported by a hierarchically-organized series of anatomically-distinguishable cortical areas, called the ventral visual stream. Early visual areas, such as primary visual cortex (V1), capture low-level features including edges and center-surround patterns^1,2^. Neural population responses in the highest ventral visual area, inferior temporal (IT) cortex, contain linearly separable information about object category that is robust to significant variations present in natural images^3,4,5^. Mid-level visual areas such as V2, V3, and V4 are less well understood, but appear to perform intermediate computations between simple edges and complex objects, correlating with sequentially increasing receptive field size^6,7,8,9,10,11,12,13,14^.

Recently, significant progress has been achieved in approximating the function of the adult primate ventral visual stream through using Deep Convolutional Neural Networks (DCNNs), a class of models directly inspired by many of these neurophysiological observations^15,16^. After being trained to learn image categorization tasks from large numbers of hand-labelled images, DCNNs have yielded the most quantitatively accurate predictive models of image-evoked population responses in early, intermediate, and higher cortical areas within the ventral visual stream^17,18,19^. The behavioral error patterns generated by these networks have also proven consistent with those of humans and non-human primates^20^. Notably, such networks are *not* directly optimized to fit neural data, but rather to solve ecologically-meaningful tasks such as object recognition. Strong neural and behavioral predictivity just “falls out” as a consequence of the high-level functional and structural assumptions constraining the networks’ optimization. Similar task-based neural network optimization approaches have led to successes in modeling the human auditory cortex^21^ and aspects of motor cortex^22^. These results suggest that the principle of “goal-driven modeling”^23^ may have general utility for modeling sensorimotor systems.

Though this progress at the intersection of deep learning and computational neuroscience is intriguing, there is a fundamental problem confronting the approach: typical neural network models of the ventral stream are built via supervised training methods involving huge numbers of semantic labels. In particular, today’s best models of visual cortex are trained on ImageNet, a dataset that contains millions of category labeled images organized into thousands of categories^24^. Viewed as a technical tool for machine learning, massive supervision can be acceptable, although it limits the purview of the method to situations with large existing labelled datasets. As a real model of biological development and learning, such supervision is highly implausible, since human infants and non-human primates simply do not receive millions of category labels during development^25,26,27^. Put another way, today’s heavily supervised neural-network based theories of cortical function may effectively proxy aspects of the real behavioral constraints on cortical systems, and thus be predictively accurate for adult cortical neural representations, but they cannot provide a correct explanation of how such representations are learned in the first place. Identifying *unsupervised* learning procedures that achieve good performance on challenging sensory tasks, and effective predictions of neural responses in visual cortex, would thus fill a major explanatory gap.

## 1 The recent emergence of high-performing unsupervised neu-ral networks

Substantial effort has been devoted to unsupervised learning algorithms over several decades, with the goal of learning task-general representations from natural statistics without high-level labelling. Early progress came from sparse autoencoding, which, when trained in shallow network architectures on natural images, produces edge-detector-like response patterns resembling some primate V1 neurons^28^. However, when applied to deeper networks, such methods have not been shown to produce representations that transfer well to high-level visual tasks or match neural responses in intermediate or higher visual cortex. More recent versions of autoencoders have utilized variational objective functions^29^, with improved task transfer performance. Unsupervised learning is also addressed in the predictive coding framework^30^, where networks learning to predict temporal or spatial successors to their inputs have achieved better task transfer^31^ and improved biological similarity^32^. Self-supervised methods, such as image colorization^33^, image context prediction^34^, and surface-normals/depth estimation^35^, have also exhibited improved task transfer.

In the last two years, a new family of unsupervised algorithms has emerged with substantially improved transfer performance, approaching that of fully-supervised networks^36,37,38,39,40,41^ These methods, which we term *contrastive embedding* objectives, include Contrastive Multiview Coding^36^ (CMC), Instance Recognition^37^ (IR), Momentum Contrast^39^ (MoCo), Simple Contrastive Learning of Representation^40^ (SimCLR), and Local Aggregation^38^ (LA). They optimize DCNNs to embed inputs into a lower-dimensional compact space, i.e. functions 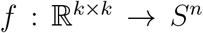, where 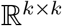 is the high-dimensional euclidean space containing *k × k* image bitmaps (with *k* ∼ 10^3^) and *S^n^* is the *n*-dimensional unit sphere (*n* ~ 128). For any input *x*, the goal is to make the embedding *f*(*x*) “unique” — that is, far away in the embedding space from other stimuli, but close to different views of the original stimulus. All methods in this family share this optimization goal, though they differ significantly in the exact implementations to achieve this uniqueness and the definition of what different “views” are. In the Local Aggregation method (Fig. 1a), for example, uniqueness is encouraged by minimizing the distance to “close” embedding points and maximizing the distance to the “further” points for each input (Fig. 1**a-b**). Through this optimization, LA explicitly seeks to create features that **generically** reflect *any* reliable natural statistic distinguishing between sets of inputs (Fig. 1**c-d**). Due to this genericity, DCNNs trained with LA create in higher network layers more and more subtle (but reliable) features for capturing whatever natural correlations are present, thus better supporting any specific high-level visual task that implicitly relies on creating distance-based boundaries in that feature space (e.g. object recognition).

**Figure 1:**
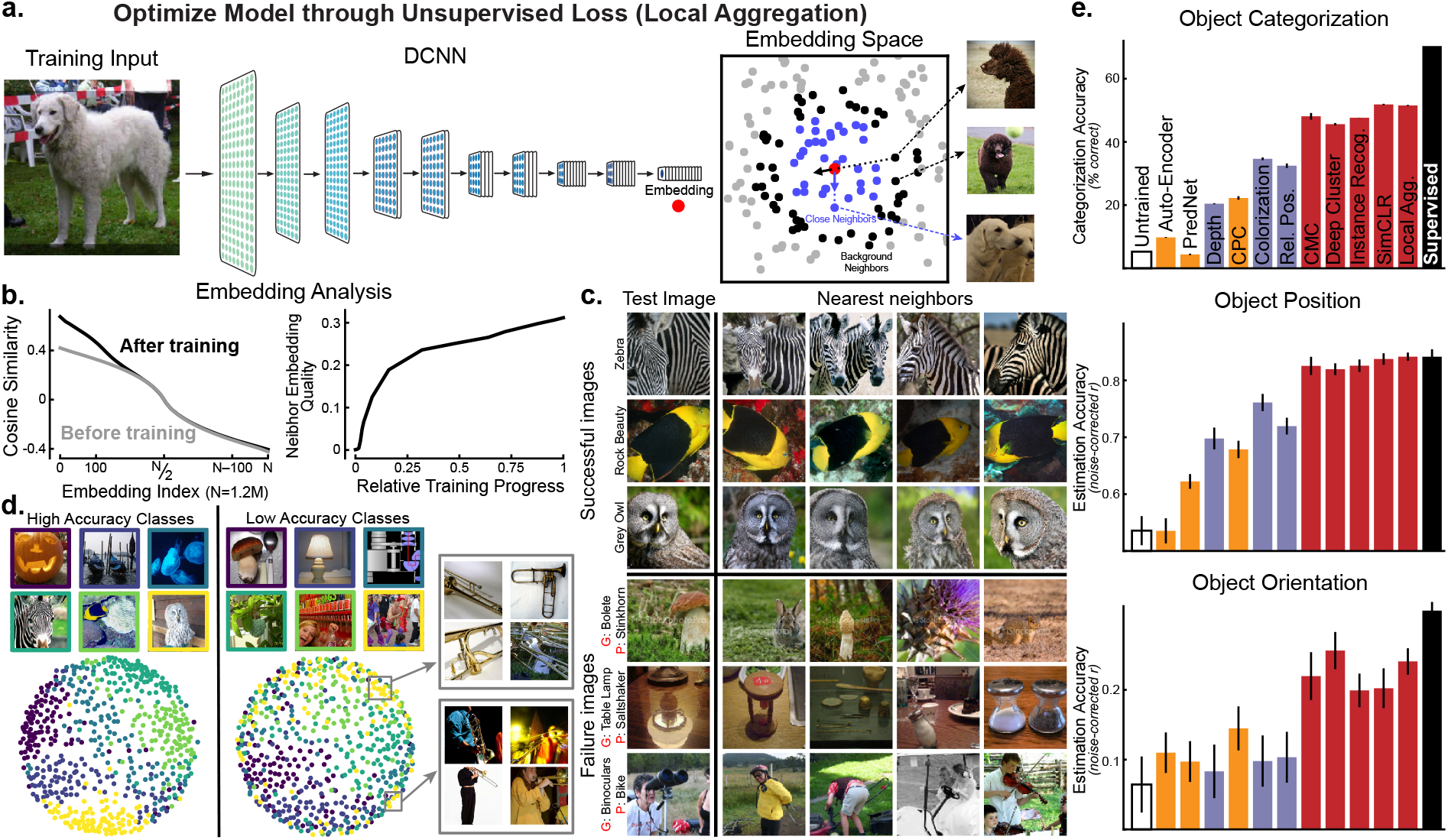
Improved representations from unsupervised neural networks based on deep contrastive embeddings. (**a**) Schematic for one high-performing deep contrastive embedding method, the Local Aggregation (LA) algorithm^38^. In LA, all images were embedded into a lower dimensional space by a DCNN, which was optimized to minimize the distance to “close” embedding points (blue dots) and to maximize the distance to the “further” points (black dots) for the current input (red dot). (**b**) (Left) Change in the embedding distribution before and after training. For each image, cosine similarities to others were computed and ranked; the ranked similarities were then averaged across all images. This metric indicates that the optimization encourages local clustering in the space, without aggregating everything. (Right) Average neighbor embedding “quality” as training progresses. Neighbor embedding quality was defined as fraction of 10 closest neighbors of the same ImageNet class label (not used in training). (**c**) Top-four closest images in the embedding space. Top three rows show the images that were successfully classified using a weighted K-Nearest-Neighbor (KNN) classifier in the embedding space (K=100), while bottom three rows show unsuccessfully-classified examples (G means ground truth, P means prediction). Even when uniform distance in the unsupervised embedding does not align with ImageNet class (which itself can be somewhat arbitrary given the complexity of the natural scenes in each image), nearby images in the embedding are nonetheless related in semantically meaningful ways. (**d**) Visualizations of Local Aggregation embedding space using Multi-Dimensional Scaling (MDS) method. Classes with high validation accuracy are shown in left panel and low accuracy classes are in right panel. Gray boxes show examples of images from a single class (“trombone”) that have been embedded in two distinct subclusters. (**e**) Transfer performance of unsupervised networks on three evaluation tasks: ImageNet categorization (upper), object position estimation (middle), and object orientation estimation (lower). Networks were first trained by unsupervised methods, then assessed on transfer performance with supervised linear readouts from network hidden layers (see Methods). Red bars are contrastive embedding tasks. Blue bars are for self-supervised tasks. Orange bars are for predictive coding methods and Auto-Encoder. White bar is the untrained model and black bar is the model supervised on ImageNet category labels. Error bars are standard variances across three networks with different initializations.

To evaluate these unsupervised learning algorithms, we trained representatives of each type described above on images from a large-scale high-quality image dataset using a fixed network architecture that has previously been shown to achieve high task performance^42^ and neural response predictivity^43^ when trained in a supervised manner. We found that these unsupervised representations achieved categorization transfer performance in line with previously reported results, validating the soundness of our implementations (Fig 1**e**, upper). Contrastive embedding objectives (red bars in Fig 1**e**) showed significantly better transfer than self-supervised tasks (blue bars), predictive coding methods (orange bars), and autoencoders. We also evaluated these representations on transfer to a variety of other object-centric visual tasks independent of object category, including object position localization and pose estimation (Fig 1**e**, middle and lower), finding that contrastive embedding objectives also achieved better transfer performance on these tasks. These results suggest that such unsupervised networks have achieved a general improvement in the quality of the visual representation.

## 2 Recent unsupervised models capture neural responses through-out ventral visual cortex

To determine whether the improvement of unsupervised methods on task transfer performance translates to better neural predictivity, we fit a regularized linear regression model from network activations of each unsupervised model to neural responses collected from array electrophysiology experiments in the macaque ventral visual pathway (Fig. 2**a**). Comparison to V1 was made using neural data collected by Cadena et al.^19^ and comparison to V4 and IT was made using data collected by Majaj et al.^3^ (see Methods for more details). Following techniques previously used to compare supervised networks to neural response patterns^17,44^, we compared each neural network separately to the neural data, and reported the noise-corrected correlation between model and neural responses across held-out images, for the best-predicting layer for each model (Fig. 2**b**). We also compared to an untrained model, which represents an architecture-only base-line. Over all, we found that the unsupervised methods that had higher transfer performance to categorization predict neural responses substantially better than less-performant unsupervised methods. All unsupervised methods were significantly better than the untrained baseline at predicting responses in early visual cortical area V1, but none were statistically distinguishable from the category-supervised model on this metric. In contrast, only a subset of methods achieve parity with the supervised model in predictions of responses in intermediate cortical area V4. (Interestingly, the deep autoencoder is not better than the untrained model on this metric and both are widely separated from the other trained models.) For IT cortex at the top of the ventral pathway, only the best-performing contrastive embedding methods achieve neural prediction parity with supervised models. Among these methods, we found that the Local Aggregation model, which has recently been shown to achieve state-of-the-art unsupervised visual recognition transfer performance^38^, also achieves the best neural predictivity. In fact, LA exhibits somewhat better V4 predictivity (two-tailed paired t-test over neurons, p=0.0087) than, as well as comparable V1 (two-tailed paired t-test over neurons, p=0.98) and IT (two-tailed paired t-test over neurons, p=0.32) predictivity to, its supervised counterpart (see Extended Data Fig. 1 for details and other t-test results). To ensure that our results are not specific to the chosen neural network architecture, we also evaluated several alternative architectures and found qualitatively similar results (Extended Data Fig. 7).

**Figure 2:**
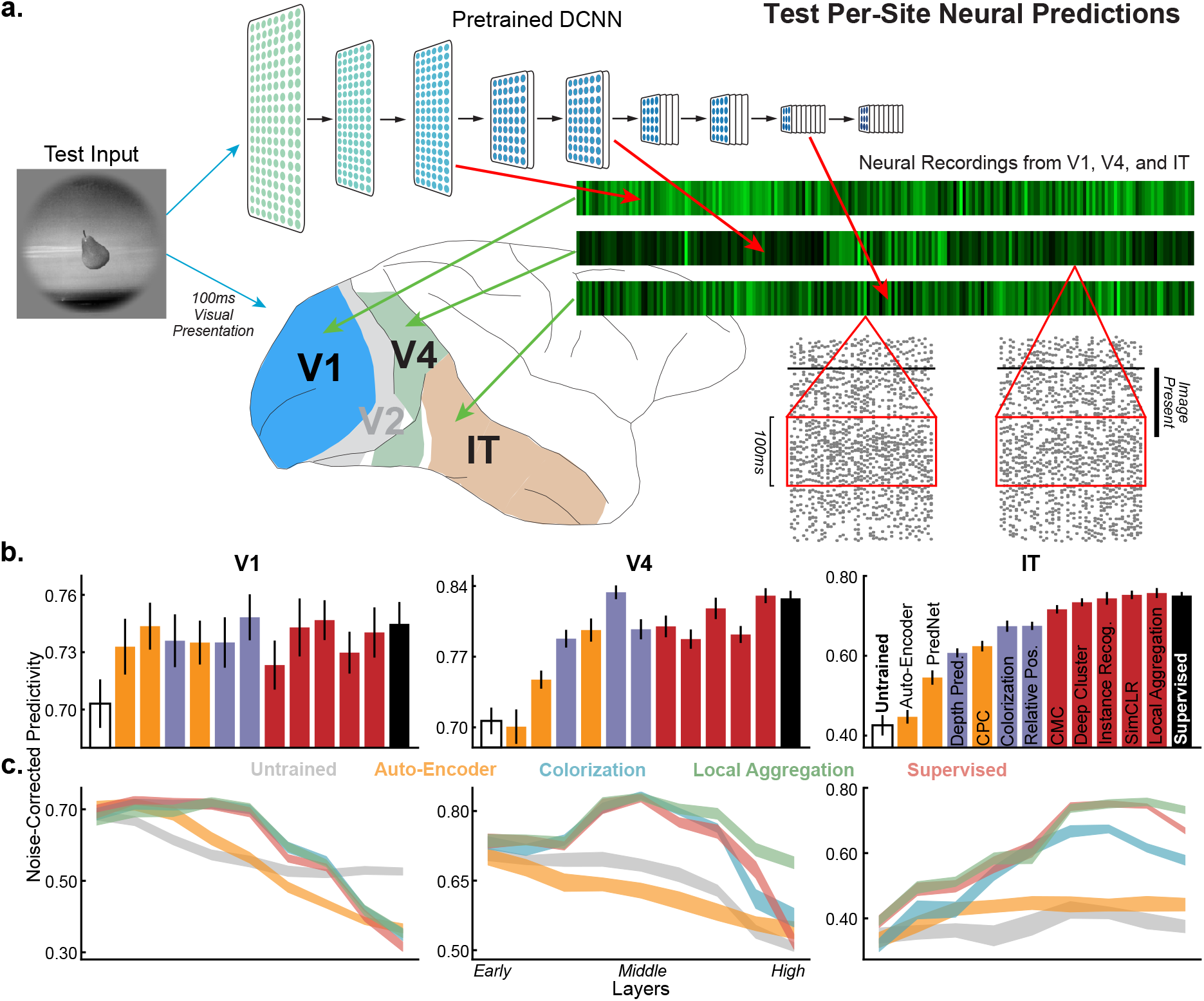
Quantifying similarity of unsupervised neural networks to visual cortex data. (**a**) After being trained with unsupervised objectives, networks were run on all stimuli for which neural responses were collected. Network unit activations from each convolutional layer were then used to predict the V1, V4, and IT neural responses with regularized linear regression^44^. For each neuron, the Pearson correlation between the predicted responses and the recorded responses was computed on held-out validation images, and then corrected by the noise ceiling of that neuron (see Methods). The median of the noise-corrected correlations across neurons for each of several cortical brain areas was then reported. (**b**) Neural predictivity of the most-predictive neural network layer. Error bars represent bootstrapped standard errors across neurons and model initializations (see Methods). Predictivity of untrained and supervised categorization networks represents negative and positive controls respectively. (**c**) Neural predictivity for each brain area from all network layers, for several representative unsupervised networks, including Auto-Encoder, Colorization, and Local Aggregation.

To further quantify the match between the computational models and brain data, we also investigated which layers of DCNNs best match which cortical brain areas (Fig. 2**c** and Extended Data Fig. 2). We found that the deep contrastive embedding models also evidence the correct model-layer-to-brain-area correspondence, with early-layer representations best predicting V1 neural responses, mid-layer representations best predicting V4 neural responses, and higher-layer representations best predicting IT neural responses. In contrast, unsupervised models with lower task performance and neural predictivity exhibit less accurate model-brain correspondence, while the untrained baseline does not show the correct correspondence at all. This conclusion is consistent across multiple quantitative metrics of mapping consistency including optimal layer match (Fig. 2**c** and Extended Data Fig. 2) as well as best predicted layer ratio metric (Extended Data Fig. 3).

In addition to the quantitative metrics described above, we also sought to qualitatively assess models. DCNNs trained with different unsupervised loss functions exhibit first layer filters with Gabor wavelet-like tuning curves like those observed in V1 data, consistent with their good neural predictivity for V1 neurons (Extended Data Fig. 4). The LA-based model, like the category-supervised model, also exhibited color-opponent center-surround units consistent with empirical observations^45,46^. Additionally, we examined optimal stimuli driving neurons in intermediate and higher model layers using techniques similar to those used in recent model-driven electrophysiology experiments^47^. Consistent with qualitative descriptions of receptive fields in the literature on V4 cortex^14^, we found that unsupervised models with good quantitative match to V4 data exhibit complex textural patterns as optimal stimuli for their most V4-like layers (Extended Data Fig. 5). In contrast, the optimal stimuli driving neurons in the most IT-like model layers appear to contain fragments of semantically-identifiable objects and scenes and large-scale organization (Extended Data Fig. 6), echoing qualitative neurophysiological findings about IT neurons^48^.

## 3 Deep constrastive learning can leverage noisy real-world video datastreams

Although we have shown that deep contrastive embedding models learn ventral-stream-like representations without using semantic labels, the underlying set of images used to train these networks — the ImageNet dataset — diverges significantly from real biological datastreams. For example, ImageNet contains single images of a large number of distinct instances of objects in each category, presented cleanly from sterotypical angles. In contrast, real human infants receive images from a much smaller set of object instances than ImageNet, viewed under much noisier conditions^49^. Moreover, ImageNet consists of statistically independent static frames, while infants receive a continuous stream of temporally correlated inputs^50^. A better proxy of the real infant datastream is represented by the recently-released SAYCam^51^ dataset, which contains head-mounted video camera data from three children (about 2 hours/week spanning ages 6-32 months) (Fig. 3**b**).

**Figure 3:**
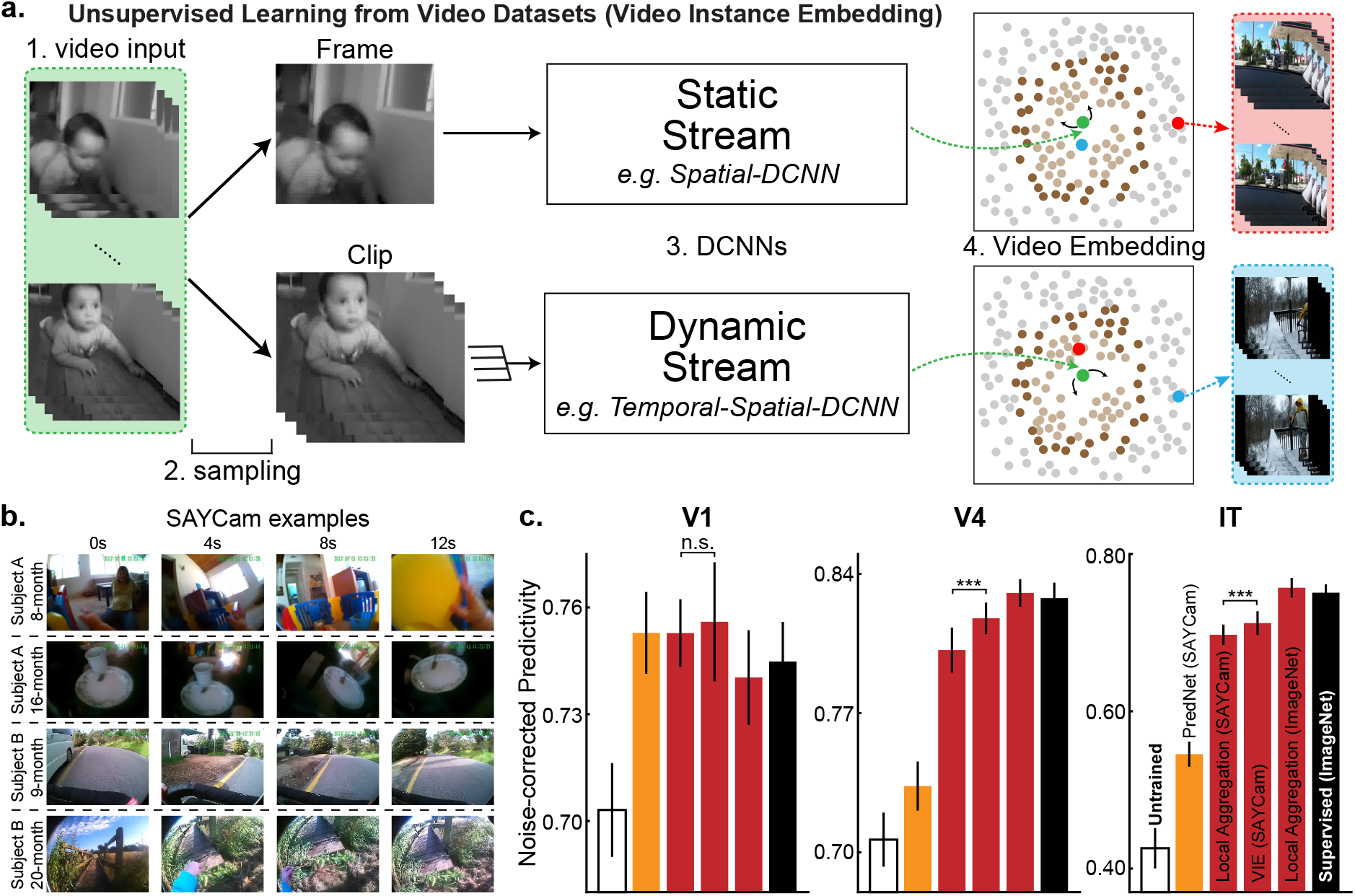
Learning from real-world developmental datastreams. (**a**) Schematic for Video Instance Embedding (VIE) method. Frames were sampled into sequences of varying lengths and temporal densities. They were then embedded into lower-dimensional space using static (single image) or dynamic (multi-image) pathways. These pathways were optimized to aggregate the resulting embeddings and their “close” neighbors (light brown points) and to separate the resulting embeddings and their “further” neighbors (dark brown points). (**b**) Examples from the SAYCam dataset^51^, which was collected by headmounted cameras on infants for two hours each week between ages 6-36 months. (**c**) Neural predictivity for models trained on SAYCam and ImageNet. “n.s.” means that the two-tailed paired t-test is not significant. “***” means highly significant t-test result (p=2.9 × 10^−9^ for V4 and p=1.9 × 10^−5^ for IT). Error bars represent bootstrapped standard errors across neurons and model initializations.

To test whether deep contrastive unsupervised learning is sufficiently robust to handle noisy and limited real-world videostreams such as SAYCam, we implemented the Video Instanace Embedding (VIE) algorithm, a recent extension of LA to video that achieves state-of-the-art results on a variety of dynamic visual tasks, such as action recognition^52^ (Fig. 3**a**). Representations learned by VIE on videos from SAYCam proved highly robust, approaching the neural predictivity of those trained on ImageNet (Fig. 3**c**). The temporally-aware VIE-trained representation was significantly (though modestly) better than a purely static network trained with LA on SAYCam frames, while both were very substantially better than PredNet, a recent biologically-inspired implementation of predictive coding^32^. These results suggest that deep spatiotemporal contrastive learning can take advantage of noisy and limited natural datastreams to achieve primate-level representation learning. A small but statistical significant gap between the SAYCam-trained and ImageNet-trained networks remains, possibly due either to limitations in the dataset (SAYCam was recorded for only two hours/week, representing a small fraction of the visual data infants actually receive) or in VIE itself.

## 4 Partial supervision improves behavioral consistency

While infants and non-human primates do not receive large numbers of semantic labels during development, it is likely that they do effectively receive at least some labels, either from parental instruction or through environmental reward signals. For human infants, object labels are provided by parents from birth onwards, but the earliest evidence for comprehension of any labels is at roughly 6-9 months of age^25^, and comprehension of most common object labels is low for many months thereafter^26^. However, visual learning begins significantly earlier at, and indeed before, birth^53^. This observation suggests that a period of what might be characterized as purely unsupervised early visual learning could be followed by a period of learning partially from labels. To capture this idea, we turned to semi-supervised learning, which seeks to leverage small numbers of labelled datapoints in the context of large amounts of unlabelled data. As with unsupervised learning, the power of semi-supervised learning algorithms has developed dramatically in recent years, benefiting from advances in understanding of neural network architectures and loss functions. The state-of-the-art existing semi-supervised learning algorithm, Local Label Propagation^54^ (LLP), builds directly on the contrastive embedding methods. Like those methods, LLP embeds datapoints into a compact embedding space and seeks to optimize a particular property of the data distribution across stimuli, but additionally takes into account the embedding properties of sparse labelled data (Fig. 4**a**). Here, we implemented both LLP and an alternative semi-supervised learning algorithm, the Mean Teacher^55^ (MT) (Extended Data Fig. 9). As precise estimates of the number of object labels available to children and the proportion of these that unambiguously label a specific object do not exist, we trained both semi-supervised models on the ImageNet dataset with 1.2M unlabeled and a range of supervision fractions, corresponding to different estimates of the number of the object speech-vision copresentations infants perceive and comprehend within the first year of life^27^. We also implemented a simple few-label control, in which standard supervision was performed using only the labeled datapoints.

**Figure 4:**
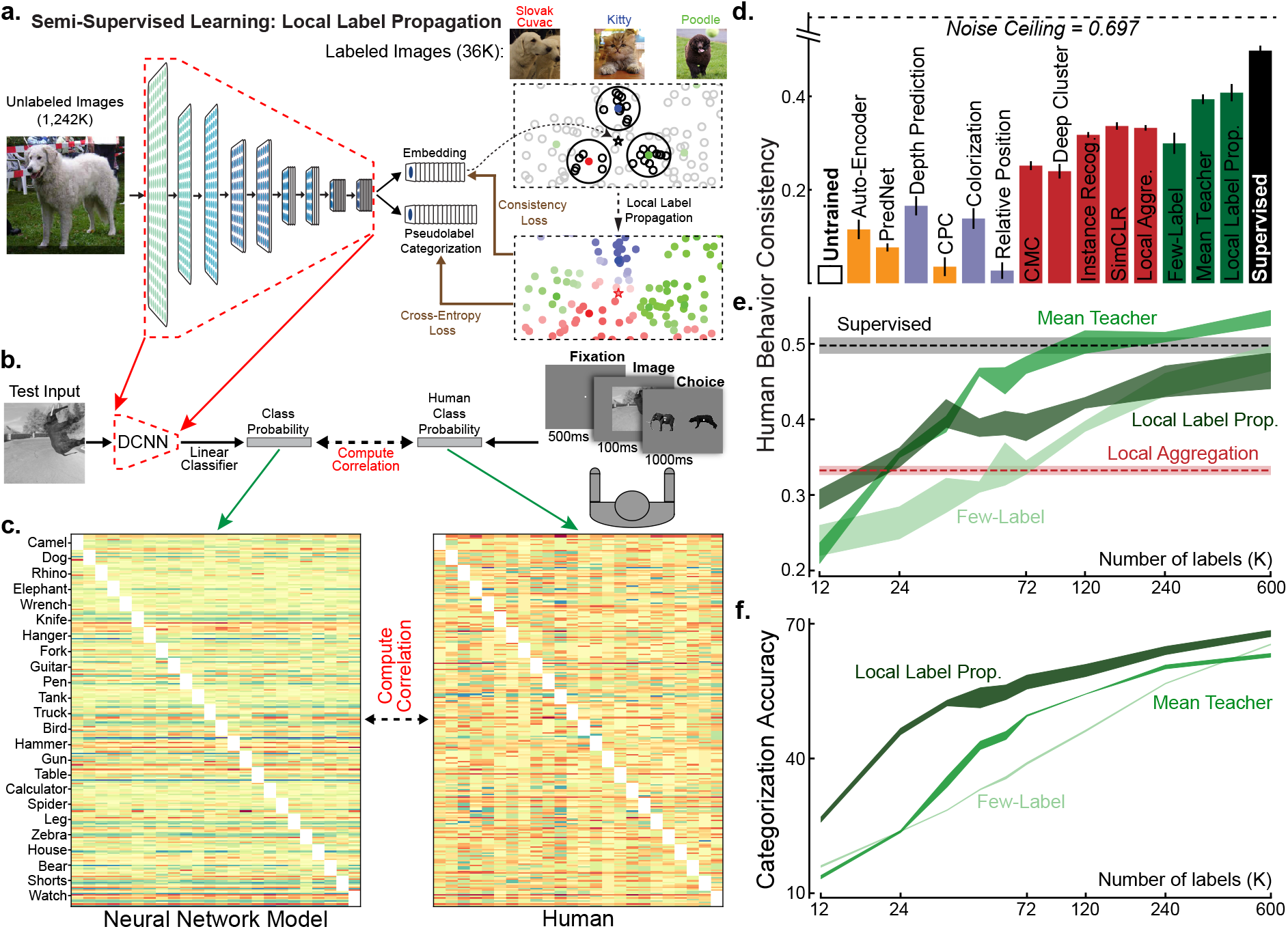
Behavioral consistency and semi-supervised learning. (**a**) In the Local Label Propagation (LLP) method^54^, DCNNs generated an embedding and a category prediction for each example. The embedding (*) of an unlabeled input was used to infer its pseudolabel considering its labeled neighbors (colored points) with voting weights determined by their distances from * and their local density (the highlighted areas). DCNNs were then optimized with per-example confidence weightings (color brightness) so that its category prediction matched the pseudo-label, while its embedding was attracted toward the embeddings sharing the same pseudo-labels and repelled by the others. (**b**) To measure behavioral consistency, we trained linear classifiers from each model’s penultimate layer on a set of images from 24 classes^20,43^. The resulting image-by-category confusion matrix was compared to data from humans performing the same alternative forced choice task, where each trial started with a 500ms fixation point, presented the image for 100ms, and required the subject to choose from the true and another distractor category shown for 1000ms^20,43^. We report the Pearson correlation corrected by the noise ceiling. (**c**) Example confusion matrixes of human subjects and model. Each category had ten images as the test images for computing the confusion matrixes. (**d**) Behavioral consistency of DCNNs trained by different tasks. Green bars are for semi-supervised models trained with 36K labels. “Few-Label” represents a ResNet-18 trained on ImageNet with only 36K images labeled, the same amount of labels used by MT and LLP models. Error bars are standard variances across three networks with different initializations. (**e, f**) Behavioral consistency (**e**) and categorization accuracy in percentage (**f**) of semi-supervised models trained with differing numbers of labels.

For each trained model, we then compared their object recognition error patterns to those in human and primates, following the methods of Rajalingham et al. ^20^, who show that category-supervised DCNNs exhibit error patterns with improved consistency to those measured in humans. We first extracted “behavior” from DCNNs by training linear classifiers from the penultimate layer of the neural network model, and measured the resulting image-by-category confusion matrix. An analogous confusion matrix was then independently measured from humans in large-scale psychophysical experiments (see Fig. 4**b**). The behavioral consistency between DCNNs and humans is quantified as the noise-corrected correlation between these confusion matrices (see Methods). We evaluated behavioral consistency both for semi-supervised models as well as the unsupervised models described above. Even using just 36K labels (corresponding to 3% supervision), both LLP and MT lead to representations that are substantially more behaviorally consistent than purely unsupervised methods, though a gap to the supervised models remains (Fig. 4**d**, see Extended Data Fig. 8 for t-test results). Interestingly, although the unsupervised LA algorithm is less consistent than either of the semi-supervised methods that feature an interaction between labelled and unlabelled data, it is more consistent than the few-label control. We find broadly similar patterns with different amounts of supervision labels (Fig. 4**e-f**). These results suggest that semi-supervised learning methods may capture a feature of real visual learning that builds on, but goes beyond, task-independent self-supervision.

## 5 Discussion

We have shown that deep contrastive unsupervised embedding methods accurately explain and predict image-evoked neural responses in multiple visual cortical areas along the primate ventral visual pathway, equaling the predictive power of supervised models. Moreover, the mapping from the layers of these unsupervised networks to corresponding cortical areas is neuroanatomically consistent, and reproduces several qualitative properties of the visual system. We have also shown that deep contrastive learning methods can take advantage of noisy and limited datasets arising from real developmental datastreams to learn strong visual representations, and that training with a semi-supervised learning objective allowing incorporation of small amounts of supervision creates networks with improved behavioral consistency with humans and non-human primates. Taken together, these results suggest that an important gap in the promising but incomplete goal-driven neural network theory of visual cortex may be close to resolution.

Contrastive embedding objectives generate image embeddings that remain invariant under certain “viewpoints” while being distinguishable from others. By minimizing these obejctives, networks effectively discover non-prespecified high-level image statistics that support reliable and generalizable distinctions^56^. This feature distinguishes the deep contrastive embedding approach from earlier unsupervised models such as autoencoders or self-supervised tasks, which optimized low-level or narrowly-defined image statistics and, as a result, learned less powerful representations. Because deep contrastive embedding methods are quite generic, and do not require the implementation of strong domain-specific priors (e.g. the presence of visual objects in 3D scenes), the application of similar methods might further understanding in other sensori-motor cortical domains where supervised neural networks have proven useful as predictors of neural responses^21,22^.

Though our results help clarify a key problem in the modeling of sensory learning, many major questions remain. Our work addresses how the visual system might develop post-natally via natural visual experience, replacing an effective but implausible learning method (heavily-supervised categorization) with one that an organism might more plausibly implement in the real world (unsupervised or semi-supervised contrastive embedding loss). However, our work does not address gaps in understanding of either the architecture class of the neural network or the low-level mechanistic learning rule.

By design, we here use the same deep feed-forward neural network architectures that have been effective for supervised learning. While such networks might be sufficient to predict temporal averages during the first volley of stimulus-evoked neural responses, they are insufficient to describe the response dynamics of real neurons^57^. Recent work has begun to integrate into neural networks analogs of the recurrences and long-range feedbacks that have been ubiquitously observed throughout the visual system, toward better modeling neural dynamics^58^. This work has been in the supervised context, so a natural future direction is to connect these architectural improvements with the unsupervised objectives explored here.

As for the mechanisms of the learning rule, our work still uses standard backpropagation for optimization (albeit with unsupervised rather than supervised objective functions). Back-propagation has several features that make it unlikely to be implementable in real organisms^59^. Historically, the question of biologically plausible unsupervised objective functions (e.g. learning targets) is intertwined with that of biologically plausible learning rules (e.g. the mechanism of error-driven update). Some specific unsupervised objective functions, such as sparse autoencoding, can be optimized with Hebbian learning rules that do not require high-dimensional error feedback^60^. However, this intertwining may be problematic, since the more effective objective functions that actually lead to powerful and neurally predictive representations do not obviously lend themselves to simple Hebbian learning. We thus suggest that these two components — optimization target and mechanism — may be decoupled, and that such decoupling might be a principle for biologically-plausible learning. This hypothesis is consistent with recent work on more biologically-plausible local learning rules that effectively implement error feedback^61^. It would be of substantial interest to build networks that use these learning rules in conjunction with unsupervised contrastive-embedding objective functions and recurrent convolutional architectures. If successful, this would represent a much more complete goal-driven deep learning theory of visual cortex.

Better training environments will also be critical. Although SAYCam is more realistic than ImageNet, there are still many important components of real developmental datastreams missing in SAYCam, including (but not limited to) the presence of *in utero* retinal waves^53^, the long period of decreased visual acuity^62^, and the lack of non-visual (e.g auditory and somatosensory) modalities that are likely to strongly self-supervise (and be self-supervised by) visual representations during development^63^. Moreover, real visual learning is likely to be at some level driven by interactive choices on the part of the organism, requiring a training environment more powerful than any static dataset can provide^64,65^.

Unsupervised deep contrastive embedding methods are more ecologically plausible than heavily supervised learning, in that it is possible to imagine a simple neural circuit by which the organism could encode the contrastive loss objective, operating just on the sensory data the organism naturally receives during post-natal development and beyond. However, it is interesting to note that the neural predictivity of the best unsupervised method only slightly surpasses that of supervised categorization models. Moreover, the detailed *pattern* of neural predictivities across units of the best unsupervised models also generally aligns with that of the supervised models (Extended Data Fig. 10). One possible explanation for these outcomes is that visual categorization is indeed a good description of the larger evolutionary constraint that the primate visual system is under, while the unsupervised algorithm is best understood as a developmental proxy for how other inaccessible representational goals might be “implemented” by the organism. Another possibility is that the neurophysiological data used in this study simply does not have the power to resolve differences between the supervised and best unsupervised models. A third possibility is that better unsupervised learning methods yet to be discovered will achieve improved neural predictivity results, substantially surpassing that of categorization models.

Ultimately, a theory of visual post-natal development should go beyond just predicting neural responses in adult animals, and also provide a model of *changes* over the time line of post-natal development. The long-term learning dynamics of any model generates trajectories of observables that could in principle be compared to similar observables measured over the course of animal development. The concept of such developmental trajectory comparison is illustrated in Extended Data Fig. 11, where we show the trajectories of observables including orientation selectivity, task performance, and an analog of neural maturation rate, over the course of *“in-silico* development.” Treating each distinct unsupervised objective function as a different hypothesis for the learning target of visual development, the comparison of these curves can be seen to successfully distinguish between the various hypotheses, even when the final “adult” state may not easily separate them. To the extent that measurements of these (or other) observables can be made over the course of biological development, it would then be possible to determine which model(s) are closest to the true developmental trajectory, or to convincingly falsify all of them. The specific observables that we measure here *in silico* may not be easily experimentally accessible, as developmental neuroscience remains technically challenging. However, our results suggest a strong motivation for turning a recent panoply of exciting technical neuroscience tools^66,67^ toward the developmental domain. In the context of model-driven experimental designs, such measurements would be of great value not only to provide insights into how visual learning proceeds, but could also inspire better unsupervised or semi-supervised learning algorithms.

## Methods

### Neural Network Training

We used ResNet-18 ^42^ without its final pooling layer and the categorization readout layer as visual backbones for all tasks except PredNet. For each task, we trained three networks with different network initializations. Most tasks were performed by adding an additional header upon the visual backbone and then training the whole network with the task-specific loss in addition to a L2-regularization loss of the network weights with a weight decay coefficient 10^−4^. Unless specified, the input image to the networks was in resolution 224 × 224 and there were two learning rate drops during training. The learning rate was dropped by 10 times after the validation performance saturates. Most training hyperparameters such as the batch size and the initial learning rate were set according to the papers of these tasks. As the Local-Aggregation task is already introduced in the main text, we only briefly describe other tasks below, of which the details can be found in their corresponding papers. For each task, we trained three networks with different initializations.

#### Auto-Encoder

The image was first projected into a 128-dimension hidden vector. An output image was then generated from this vector using a reversed ResNet-18. The Auto-Encoder loss optimized was the sum of the L2 different between the output and the input images and the L1-norm of the hidden vector multiplied by 10^−4^.

#### PredNet^68^

PredNet network was trained to predict the next frame using previous frames and a specifically-designed recurrent neural network architecture including four modules each of which has three layers. As ResNet-18 is a feedforward network, it cannot be used in PredNet. We also found that PredNet failed to train more layers added to each module. Therefore, we used the same network architecture as the original paper^68^. Later neural fitting, object position/pose estimation task, and categorization tasks were all performed using each of the twelve layers and we reported the best number across these layers. As the network architecture is very different from others, we cannot show comparable neural fitting layer trajectories.

#### Depth Prediction

A multi-layer header was added to the visual backbone to output a per-pixel depth image^35^. This depth image was then compared the ground truth normalized depth image, of which the mean was 0 and the standard deviation was 1 within one image. The L2-norm of the difference was used as the optimization loss. This task was trained on PBRNet, which is a large-scale synthetic dataset containing 0.4 million images which are physically-based rendered from 45K realistic 3D indoor scenes^69^.

#### Contrastive Predictive Coding (CPC)^31^

A grid of small crops were taken from the input image and then fed into the network to generate a grid of low-dimension embeddings. The network was then optimized to predict one embedding from its spatial neighbors using a re-current head. Although this method has “contrastive” in its name, CPC is very different from deep contrastive embedding methods as it predicts the current embedding in the context of the embeddings of all the other small crops within the same image, while for deep contrastive embedding methods, the “contrastive” usually represents the context of the embeddings of other examples.

#### Colorization^33^

The input image was first converted into Lab color space. The L channel was then used as the input to the network to predict the ab channels.

#### Relative Position^34^

Two small image crops were first chosen from a 3 × 3 grid of small image crops and then fed into the network. Their outputs were then used to predict the relative spatial position using a multi-layer header.

#### Contrastive Multiview Coding (CMC)^36^

The input image was first converted into Lab color space. Both L and ab channels were embedded into 128-dimension vectors using two different visual backbones. These two visual backbones were then optimized to push together the vectors of corresponding L and ab channels and to separate them away from the embeddings of other images. The L-ResNet18 was used to evaluate the neural predictivity and the behavior consistence metric, as the stimulus used there are all gray-scale images.

#### Deep Cluster^41^

All training images were first embedding into a lower-dimension space. These embeddings were then clustered into small clusters using the KMeans algorithm. The index of the cluster one image belonged to was used as a category label for this image to train the network. These steps were iterated to get the final network.

#### Instance Recognition^37^

The input image was embedded into a 128-dimension space and the network was then optimized to separate the current embedding from all other embeddings in this space and also to aggregate the current embedding and the running-average of embeddings of previous training examples from the same image, which are different data augmentation instances.

#### SimCLR^40^

SimCLR has the same objective as Instance Recognition, but is different from it in its architecture between visual backbone and the embedding, more data-augmentations, a different hyper-parameter, and a bigger batch-size during training. We used the official codes from the authors to train our SimCLR ResNet-18.

#### VIE^52^

We trained two pathways using VIE loss on SAYCam: static pathway with ResNet-18 and dynamic pathway with 3D-ResNet-18, which receives 16 consecutive frames and applies temporal and spatial convolutions^52^. The pretrained two pathways were concatenated across the channel dimension in each layer as the final network. When testing on static stimuli, we repeated the images for 16 times and averaged the responses of the dynamic pathway across the temporal dimension.

### Neural Predictivity Evaluation

#### Neural response dataset for V1 area

This dataset was collected by presenting stimulus to two awake and fixating macaques, where responses of 166 neurons in V1 area were collected by a linear 32-channel array^19^. The stimulus consisted of 1450 images from ImageNet and texture-like synthesized images matching the outputs of different layers of a ImageNet trained deep neural networks. The images were presented for 60ms each in one trial without blanks and centered on the population receptive field of the neurons in each session. A total of 262 neurons were isolated in 17 sessions. The response latency of these neurons is typically 40ms. Therefore, spike counts between 40-100ms were extracted and averaged across trials to get the final responses. Neurons were further selected based on whether at least 15% of their total variance could be attributed to the stimuli, which left 166 neurons.

#### Neural response dataset for V4 and IT areas

This dataset was collected by presenting stimuli to two fixating macaques, on which three arrays of electrodes were implanted with one array in area V4 and the other two arrays in area IT^3^. The stimuli were constructed by rendering one of 64 3-dimensional objects at randomly chosen position, pose, and size, on a randomly chosen naturalistic photograph as background. These objects belonged to 8 categories (tables, planes, fruits, faces, chairs, cars, boats, animals), each of which consisted of 8 unique objects. According to the scale of variances position, pose, and size are sampled from, three datasets were generated, corresponding to low, medium, and high variations. For example, low variation images had objects placed in the center of the images with a fixed position, pose, and size, while objects in high variation images are placed with a highly varied setting. There were in total 5,760 images, of which 2,560 were high variation images and 2,560 were medium variation images. These images were presented to the primates for 100ms with 100ms of gap between images. During presentation, a circular mask was applied to each image, which subtended around 8 degree of visual angle. From the three arrays, the neural responses of 168 IT neurons and 88 V4 neurons were collected^17,3^. Following the previous studies^17^, we used the averaged responses between 70-170ms after stimuli presentation as this window contained most of object category-related information. The low and medium variation images were used to select hyperparameters of neural fitting and only the prediction results on high variation images were reported.

### Downstream Task Performance Evaluation

#### ImageNet categorization task

A linear readout layer was added to the pretrained visual backbones. This layer was trained to perform the ImageNet categorization task through a cross-entropy loss. The initial learning rate was 0.01 and the training took 160 epochs. We dropped the learning rate at 40th, 80th, and 140th epochs. Each learning rate drop was by 10 times. A L2 regularization loss on the linear readout weights was used with the regularization coefficient 10^−4^. We used the same data augmentations used in previous studies^37,38^: random cropping, random horizontal flip, random color jittering, and random gray-scale transform. We reported the best categorization performance on the official ImageNet validation set throughout the training.

#### Object position estimation task

We used the same dataset, on which the V4 and IT neural data was collected. For each pretrained visual backbone, we took the spatially averaged outputs from the first pooling layer and all eight residual blocks and then regressed them to predict both the vertical and the horizontal locations of the object center in the image. We chose the fitting hyperparameters on the low and medium variation subsets and tested the fitting on the high variation subset. The Pearson correlations between the predicted and the ground truth positions were computed for all layers. For each network, we picked the best layer using the separate dataset and reported the correlation averaged across both locations.

#### Object pose estimation task

The only thing different in this task compared to position pre-diction task was that the prediction target was the z-axis (vertical axis) and the y-axis (horizontal axis) rotations.

#### Neural response fitting procedure

For the neural network response *n_o_* of one layer whose output shape is [*s_x_, s_y_, c*], we fit a spatial mask *m_s_* of shape [*s_x_, s_y_*] and a channel mask *m_c_* of shape [*c*] for each neuron to predict its response *r*. The predicted response can be written as:

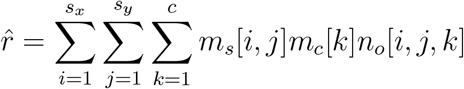

The optimized loss is then:

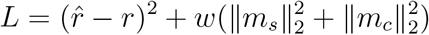

We chose *w* on low and medium variation images used in collecting V4 and IT neural responses. The weights were trained on the training split and evaluated on the validation set. The Pearson correlation was computed between 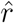 and *r* and further corrected by being divided by the square root of cross-trial neural response correlation^43^. The median value of the corrected correlations of all neurons within one cortical area was reported for one layer as its neural predictivity for this area. For each neuron, the neural predictivity is first averaged across all three networks. The error bars are the standard variances of means generated through bootstrapping from neural predictivity of all neurons for 200 times.

### Optimal Stimuli Computation

Following a previous study^70^, we optimized the 2D discrete Fourier transform of the input image to maximize the spatially averaged responses of one channel in a given layer. We used Adam optimizer with learning rate 0.05 and trained for 512 steps. In each training step, random augmentations including jitterring, scaling, and rotations were applied to the input image^70^.

### Human Behavior Consistence Metric

The behavior dataset consists of 2400 images generated by putting 24 objects in front of high-variant and independent naturalistic backgrounds^20^. For each pretrained network, a linear classifier was trained from the penultimate layer on 2160 training images to predict the category. As the number of features may be too large, we tested three dimension reduction methods and report the best consistency among them. These methods are: 1. averaging across spatial dimensions; 2. PCA projections to 1000 components using ImageNet validation images; 3. Training a 1000-dimension linear category-centric projection through adding a linear layer outputing 1000 units upon the current layer and another linear readout upon this linear layer and then training only these two added layers to do ImageNet categorization tasks. The resulted confusion matrix on 240 validation images was compared to that of human subjects, for which 1,472 human subjects were recruited from Amazon Mechanical Turk (details can be found in Rajalingham et al.^20^ and Schrimpf et al.^43^). The Pearson correlation between the matrixes was computed and then corrected by dividing it using the square root of human split-half correlation^43^.

## Data Availability

ImageNet can be downloaded from http://www.image-net.org/. SAYCam can be downloaded from https://nyu.databrary.org/volume/564. Neural data is available through the public Brain-Score repo.

## Code Availability

Our codes can be found at https://github.com/neuroailab/unsup_vvs.

## Author Contributions

C.Z., M.C.F, and D.L.K.Y. designed the experiments, C.Z. and S.Y. conducted the experiments, and C.Z. analysed the data. A.N. provided technical advice on neural network training. M.S. and J.J.D. provided technical advice on neural predictivity metrics. C.Z. and D.L.K.Y. interpreted the data and wrote the paper.

## Competing Interest Declaration

The authors declare no competing interests.

## Extended Data

**Figure 1:**
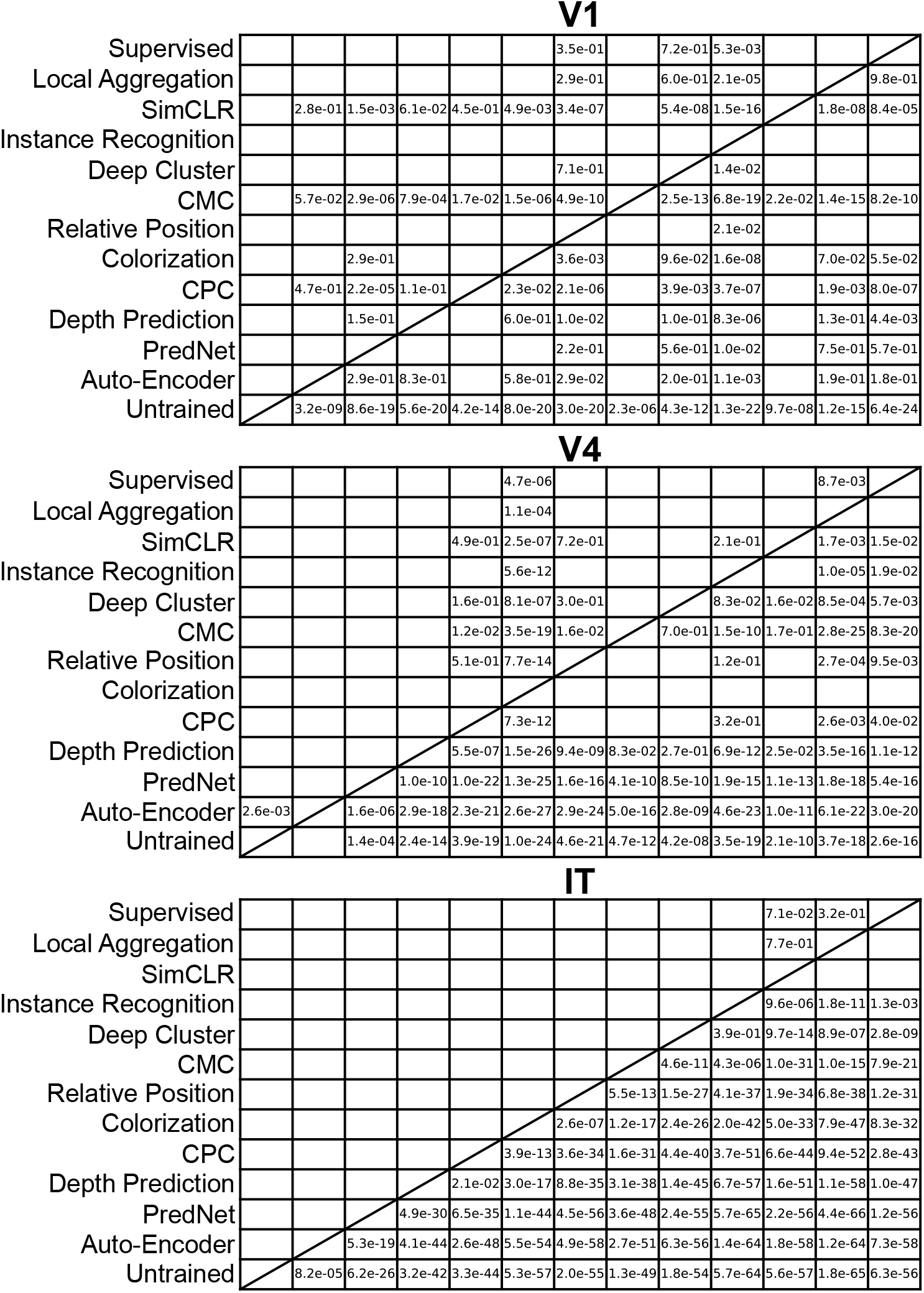
T-test results of neural predictivity results. If the neural predictivity of the method in the i-th row is smaller than that of the method in the j-th column, the number in the i-th row and j-th column then means the paired and two-tailed t-test p-value bewteen two methods. For V1, the degree of freedom is 165. For V4, the degree of freedom is 87. For IT, the degree of freedom is 167.

**Figure 2:**
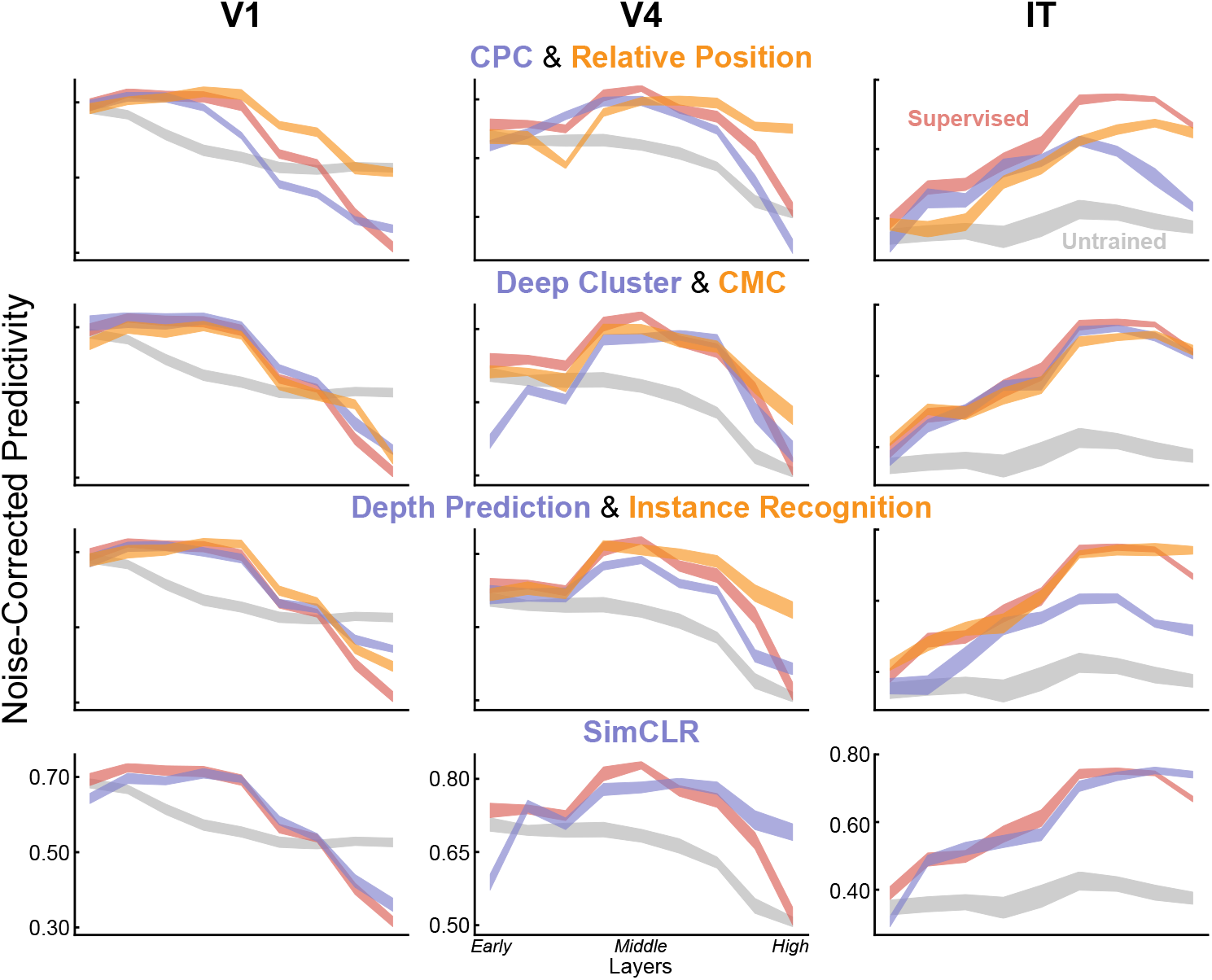
Neural predictivity of DCNNs across layers.

**Figure 3:**
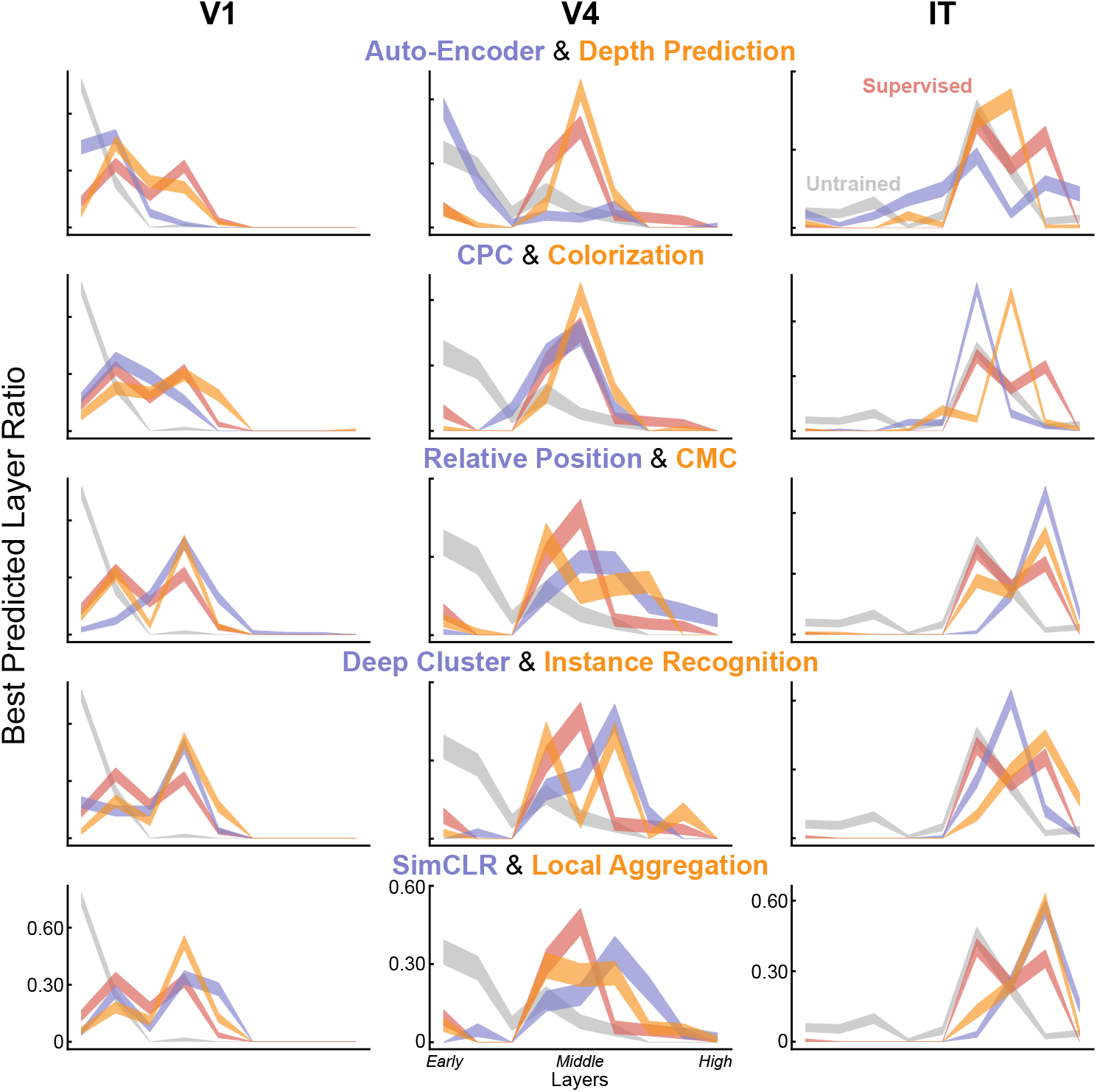
Best predicted layer ratio for neurons. For each DCNN layer, we compute the number of neurons that are best predicted by this layer and then divide this number by the number of neurons of this area.

**Figure 4:**
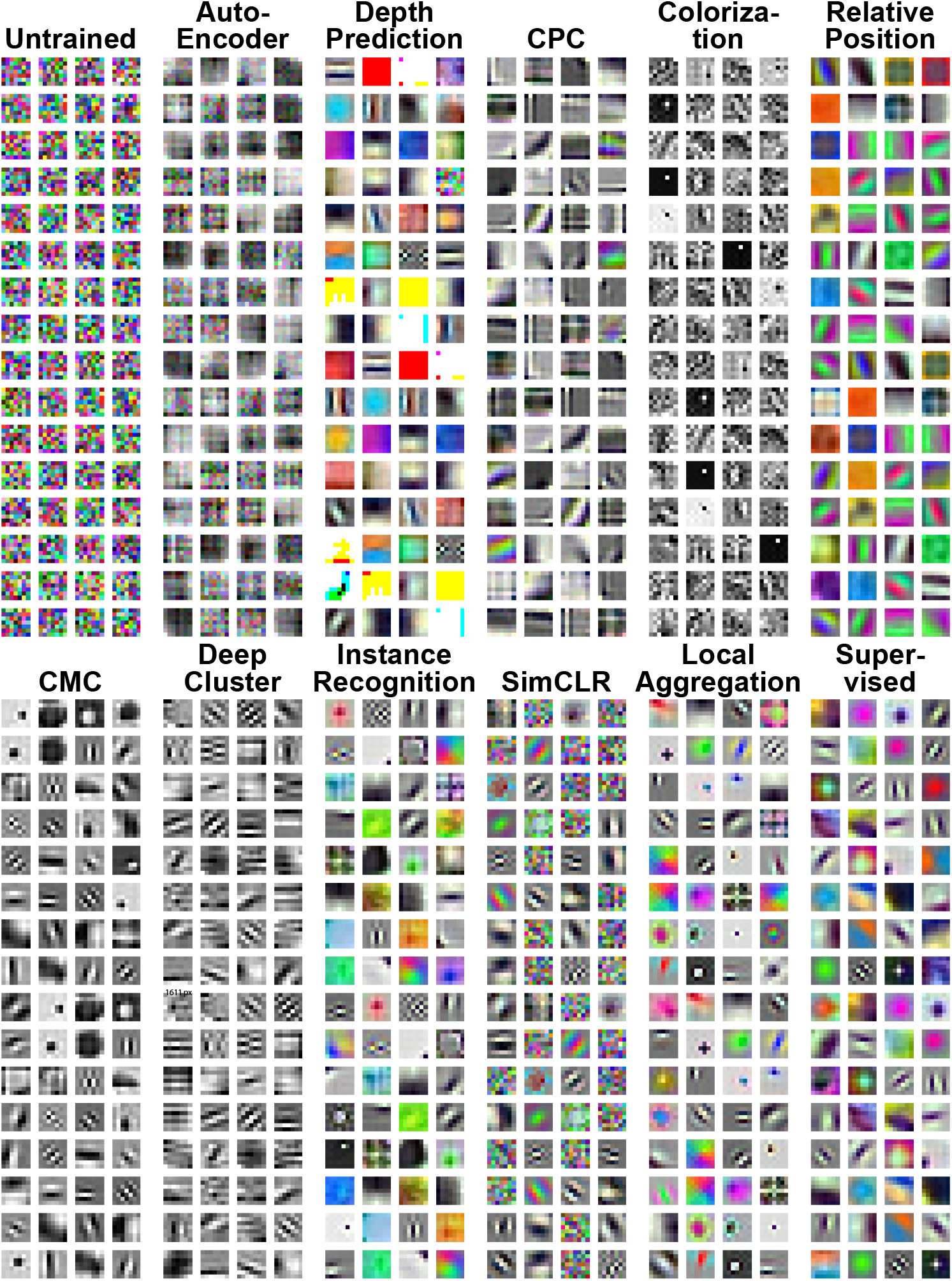
First layer filters of DCNNs.

**Figure 5:**
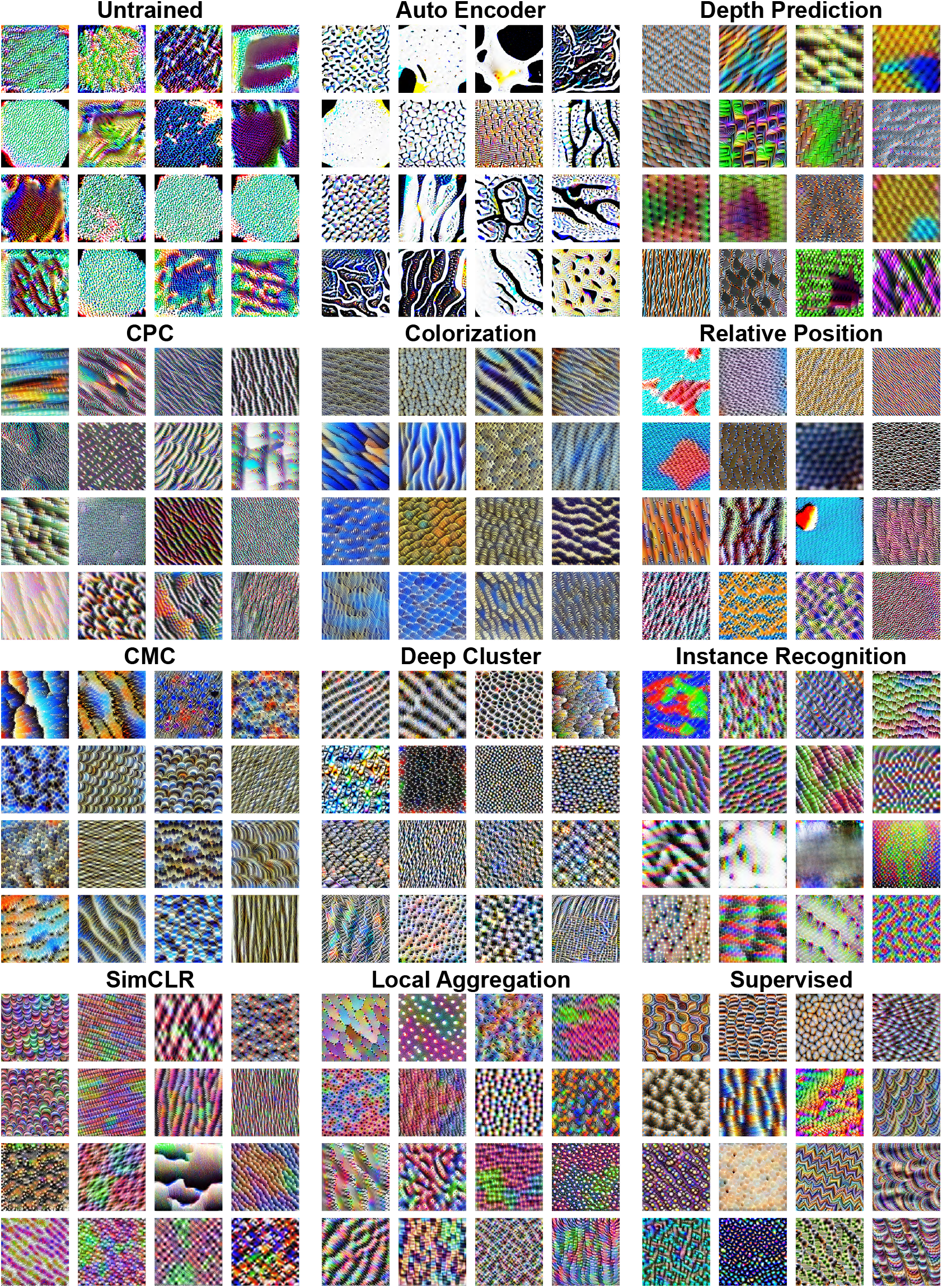
Optimal stimuli for intermediate layers of DCNNs.

**Figure 6:**
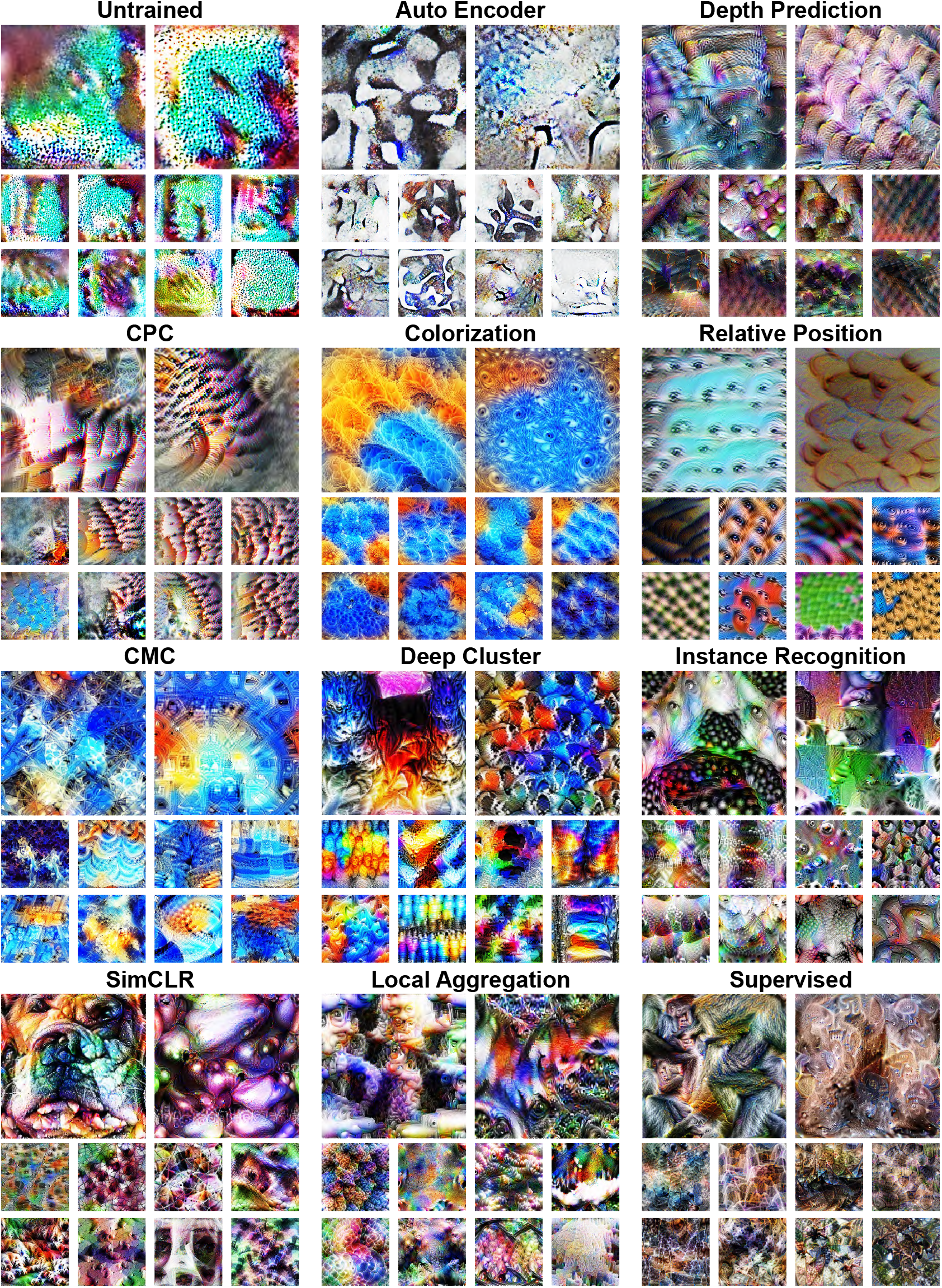
Optimal stimuli for high layers of DCNNs. Best viewed when scaled.

**Figure 7:**
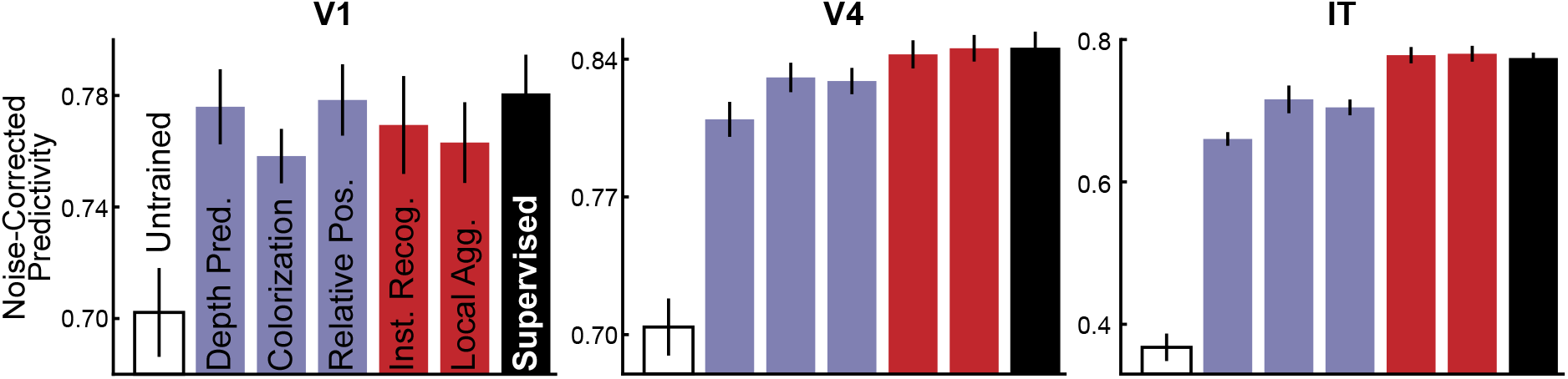
Neural predictivity results of ResNet-50 models.

**Figure 8:**
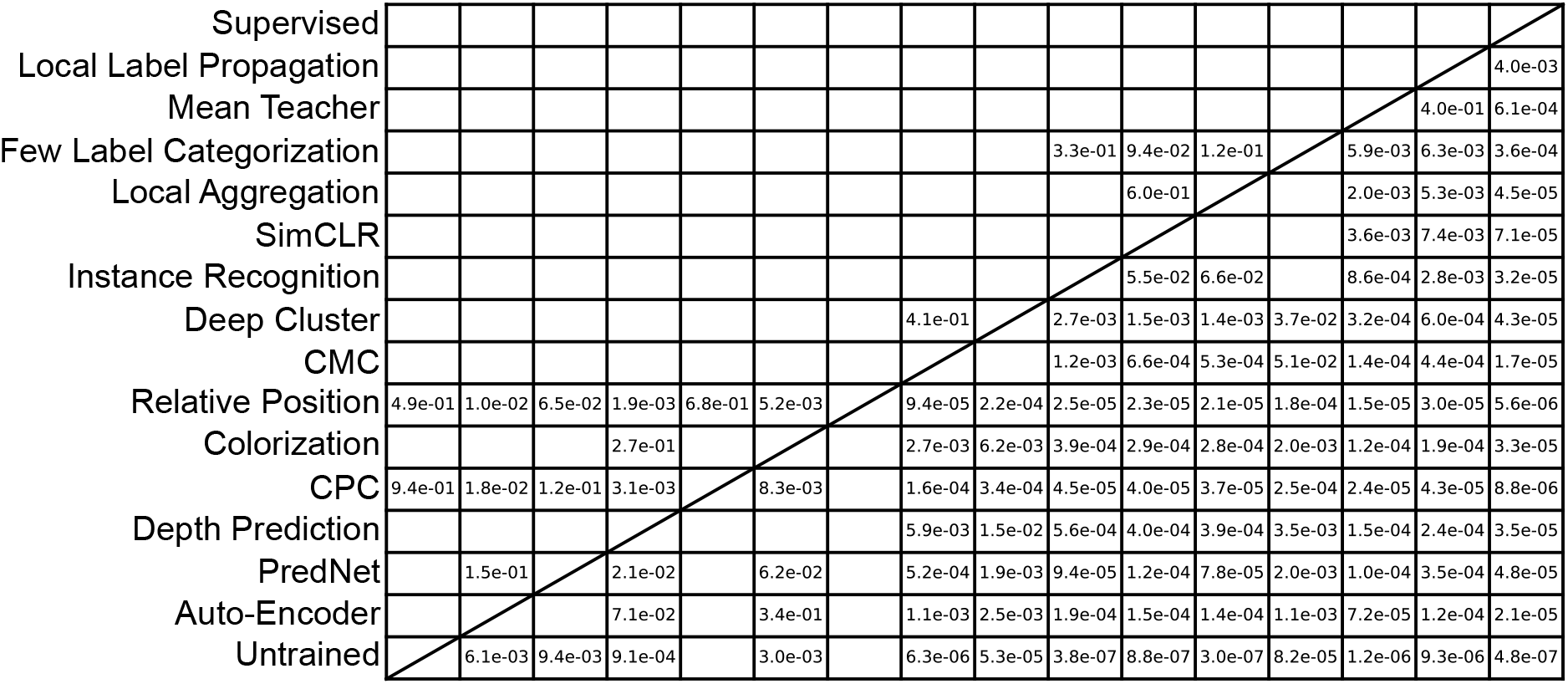
T-test results of human behavior consistency results (Fig 4**d**). If the human behavior consistency of the method in the i-th row is smaller than that of the method in the j-th column, the number in the i-th row and j-th column then means the unpaired and two-tailed t-test p-value bewteen two methods. The degree of freedom is 2, as each model has three networks of different initializations.

**Figure 9:**
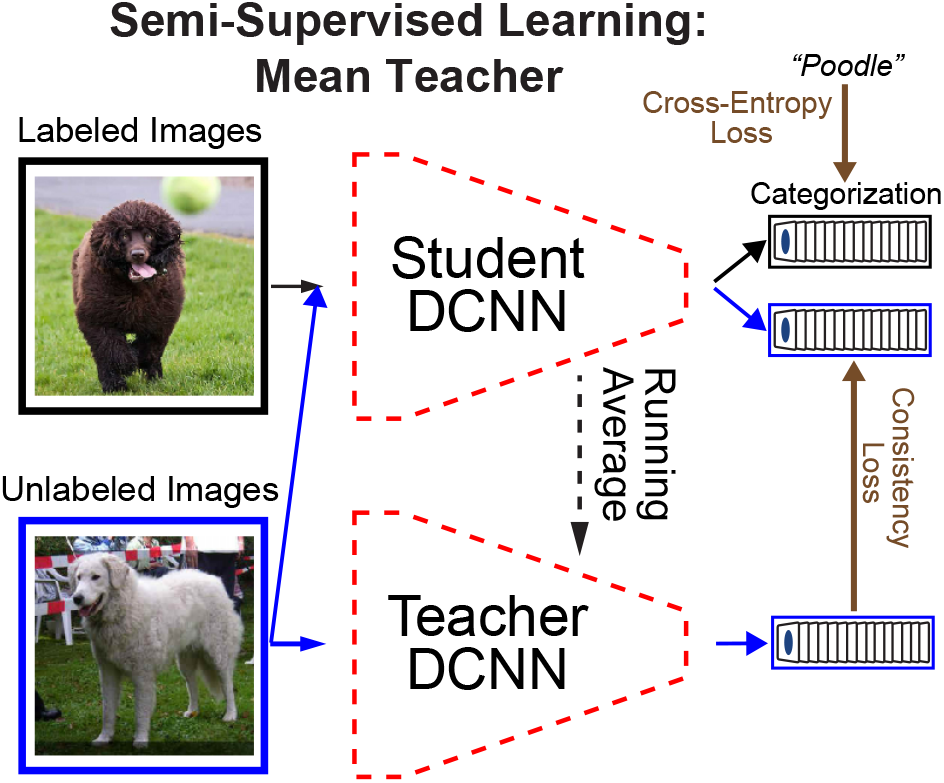
Schematic for the Mean Teacher (MT) method. In addition to the optimized “student DCNN”, MT maintained a “teacher DCNN”, whose weights were running averages of the student DCNN. The loss was the sum of the categorization loss on labeled images and the consistency loss between two DCNNs on unlabeled images.

**Figure 10:**
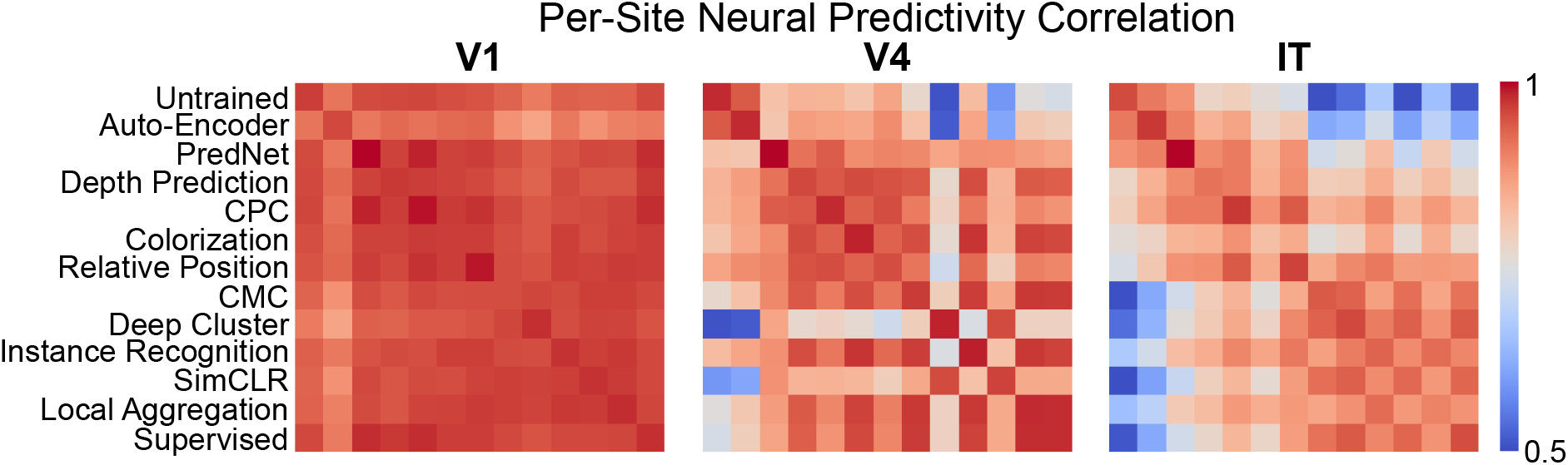
Correlation of per-site neural predictivity results of different DCNNs.

**Figure 11:**
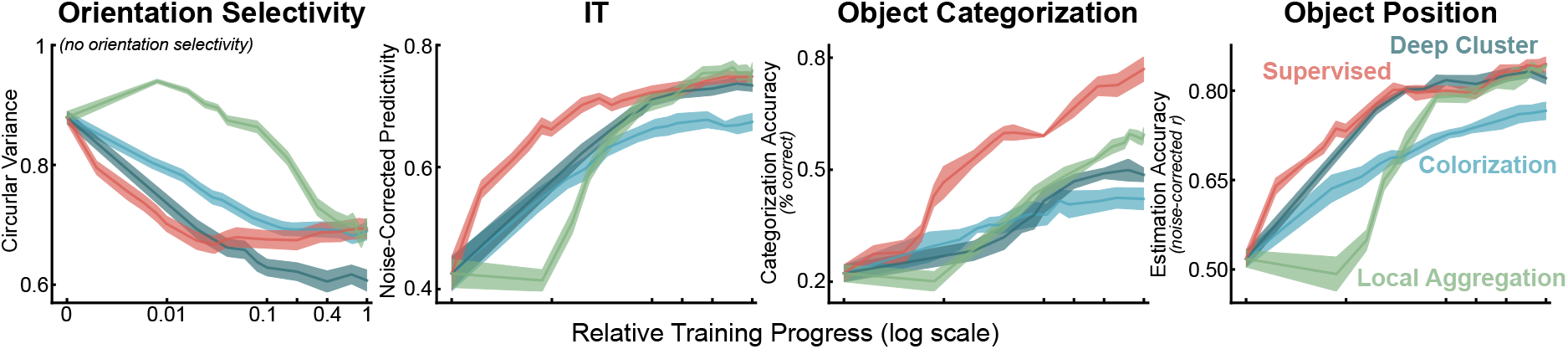
Training trajectories for (left to right): orientation selectivity measured by circular variance on early layer of DCNNs, IT neural predictivity of best layer, object categorization, object position predicting performance.

## Notes

### Competing Interest Statement

The authors have declared no competing interest.

## References

[1] Carandini, M. et al. Do we know what the early visual system does? J Neurosci 25, 10577–97 (2005).

[2] Movshon, J. A., Thompson, I. D. & Tolhurst, D. J. Spatial summation in the receptive fields of simple cells in the cat’s striate cortex. The Journal of physiology 283, 53–77 (1978).

[3] Majaj, N. J., Hong, H., Solomon, E. A. & DiCarlo, J. J. Simple learned weighted sums of inferior temporal neuronal firing rates accurately predict human core object recognition performance. Journal of Neuroscience 35, 13402–13418 (2015).

[4] Yamane, Y., Carlson, E. T., Bowman, K. C., Wang, Z. & Connor, C. E. A neural code for three-dimensional object shape in macaque inferotemporal cortex. Nat Neurosci (2008).

[5] Hung, C. P., Kreiman, G., Poggio, T. & Dicarlo, J. J. Fast readout of object identity from macaque inferior temporal cortex. Science 310, 863–866 (2005).

[6] Freeman, J. & Simoncelli, E. Metamers of the ventral stream. Nature Neuroscience 14, 1195–1201 (2011).

[7] DiCarlo, J. J. & Cox, D. D. Untangling invariant object recognition. Trends Cogn Sci 11, 333–41 (2007).

[8] DiCarlo, J. J., Zoccolan, D. & Rust, N. C. How does the brain solve visual object recognition? Neuron 73, 415–34 (2012).

[9] Schmolesky, M. T. et al. Signal timing across the macaque visual system. J Neurophysiol 79, 3272–8 (1998).

[10] Lennie, P. & Movshon, J. A. Coding of color and form in the geniculostriate visual pathway (invited review). J Opt Soc Am A Opt Image Sci Vis 22, 2013–33 (2005).

[11] Schiller, P. Effect of lesion in visual cortical area v4 on the recognition of transformed objects. Nature 376, 342–344 (1995).

[12] Gallant, J., Connor, C., Rakshit, S., Lewis, J. & Van Essen, D. Neural responses to polar, hyperbolic, and cartesian gratings in area v4 of the macaque monkey. Journal of Neurophysiology 76, 2718–2739 (1996).

[13] Brincat, S. L. & Connor, C. E. Underlying principles of visual shape selectivity in posterior inferotemporal cortex. Nat Neurosci 7, 880–6 (2004).

[14] Yau, J. M., Pasupathy, A., Brincat, S. L. & Connor, C. E. Curvature processing dynamics in macaque area v4. Cerebral Cortex bhs004 (2012).

[15] Fukushima, K. Neocognitron: A self-organizing neural network model for a mechanism of pattern recognition unaffected by shift in position. Biol Cybernetics (1980).

[16] LeCun, Y. & Bengio, Y. Convolutional networks for images, speech, and time series. The handbook of brain theory and neural networks 255–258 (1995).

[17] Yamins, D. L. et al. Performance-optimized hierarchical models predict neural responses in higher visual cortex. Proceedings of the National Academy of Sciences 111, 8619–8624 (2014).

[18] Kriegeskorte, N. Deep neural networks: a new framework for modeling biological vision and brain information processing. Annual review of vision science 1, 417–446 (2015).

[19] Cadena, S. A. et al. Deep convolutional models improve predictions of macaque v1 responses to natural images. PLoS computational biology 15, e1006897 (2019).

[20] Rajalingham, R. et al. Large-scale, high-resolution comparison of the core visual object recognition behavior of humans, monkeys, and state-of-the-art deep artificial neural networks. Journal of Neuroscience 38, 7255–7269 (2018).

[21] Kell, A. J., Yamins, D. L., Shook, E. N., Norman-Haignere, S. V. & McDermott, J. H. A task-optimized neural network replicates human auditory behavior, predicts brain responses, and reveals a cortical processing hierarchy. Neuron 98, 630–644 (2018).

[22] Sussillo, D., Churchland, M. M., Kaufman, M. T. & Shenoy, K. V. A neural network that finds a naturalistic solution for the production of muscle activity. Nature neuroscience 18, 1025–1033 (2015).

[23] Yamins, D. L. & DiCarlo, J. J. Using goal-driven deep learning models to understand sensory cortex. Nature neuroscience 19, 356 (2016).

[24] Deng, J. et al. ImageNet: A Large-Scale Hierarchical Image Database. In IEEE CVPR (2009).

[25] Bergelson, E. & Swingley, D. At 6–9 months, human infants know the meanings of many common nouns. Proceedings of the National Academy of Sciences 109, 3253–3258 (2012).

[26] Frank, M., Braginsky, M., Marchman, V. & Yurovsky, D. Variability and consistency in early language learning: The wordbank project (2019).

[27] Bergelson, E. & Aslin, R. N. Nature and origins of the lexicon in 6-mo-olds. Proceedings of the National Academy of Sciences 114, 12916–12921 (2017).

[28] Olshausen, B. A. & Field, D. J. Sparse coding with an overcomplete basis set: A strategy employed by v1? Vision research 37, 3311–3325 (1997).

[29] Donahue, J., Krähenbühl, P. & Darrell, T. Adversarial feature learning. arXiv preprint arXiv:1605.09782 (2016).

[30] Rao, R. P. & Ballard, D. H. Predictive coding in the visual cortex: a functional interpretation of some extra-classical receptive-field effects. Nature neuroscience 2, 79–87 (1999).

[31] Oord, A. v. d., Li, Y. & Vinyals, O. Representation learning with contrastive predictive coding. arXiv preprint arXiv:1807.03748 (2018).

[32] Lotter, W., Kreiman, G. & Cox, D. A neural network trained for prediction mimics diverse features of biological neurons and perception. Nature Machine Intelligence 2, 210–219 (2020).

[33] Zhang, R., Isola, P. & Efros, A. A. Colorful image colorization. In ECCV, 649–666 (Springer, 2016).

[34] Doersch, C., Gupta, A. & Efros, A. A. Unsupervised visual representation learning by context prediction. In Proceedings of the IEEE International Conference on Computer Vision, 1422–1430 (2015).

[35] Laina, I., Rupprecht, C., Belagiannis, V., Tombari, F. & Navab, N. Deeper depth prediction with fully convolutional residual networks. In 2016 Fourth 3DV, 239–248 (IEEE, 2016).

[36] Tian, Y., Krishnan, D. & Isola, P. Contrastive multiview coding. arXiv preprint arXiv:1906.05849 (2019).

[37] Wu, Z., Xiong, Y., Yu, S. X. & Lin, D. Unsupervised feature learning via non-parametric instance discrimination. In CVPR, 3733–3742 (2018).

[38] Zhuang, C., Zhai, A. L. & Yamins, D. Local aggregation for unsupervised learning of visual embeddings. In Proceedings of the IEEE International Conference on Computer Vision, 6002–6012 (2019).

[39] He, K., Fan, H., Wu, Y., Xie, S. & Girshick, R. Momentum contrast for unsupervised visual representation learning. arXiv preprint arXiv:1911.05722 (2019).

[40] Chen, T., Kornblith, S., Norouzi, M. & Hinton, G. A simple framework for contrastive learning of visual representations. arXiv preprint arXiv:2002.05709 (2020).

[41] Caron, M., Bojanowski, P., Joulin, A. & Douze, M. Deep clustering for unsupervised learning of visual features. In ECCV, 132–149 (2018).

[42] He, K., Zhang, X., Ren, S. & Sun, J. Deep residual learning for image recognition. In Proceedings of the IEEE conference on computer vision and pattern recognition, 770–778 (2016).

[43] Schrimpf, M. et al. Brain-score: Which artificial neural network for object recognition is most brain-like? bioRxiv preprint (2018).

[44] Klindt, D., Ecker, A. S., Euler, T. & Bethge, M. Neural system identification for large populations separating what and where. In Advances in Neural Information Processing Systems, 3506–3516 (2017).

[45] Hubel, D. H. & Wiesel, T. N. Receptive fields and functional architecture of monkey striate cortex. The Journal of physiology 195, 215–243 (1968).

[46] De Valois, R. L., Yund, E. W. & Hepler, N. The orientation and direction selectivity of cells in macaque visual cortex. Vision research 22, 531–544 (1982).

[47] Bashivan, P., Kar, K. & DiCarlo, J. J. Neural population control via deep image synthesis. Science 364, eaav9436 (2019).

[48] Kobatake, E. & Tanaka, K. Neuronal selectivities to complex object-features in the ventral visual pathway of the macaque cerebral cortex. Journal of Neurophysiology 71, 856–867 (1994).

[49] Smith, L. B. & Slone, L. K. A developmental approach to machine learning? Frontiers in psychology 8, 2124 (2017).

[50] Bambach, S., Crandall, D. J., Smith, L. B. & Yu, C. An egocentric perspective on active vision and visual object learning in toddlers. In 2017 ICDL-EpiRob, 290–295 (IEEE, 2017).

[51] Sullivan, J., Mei, M., Perfors, A., Wojcik, E. H. & Frank, M. C. Saycam: A large, longitudinal audiovisual dataset recorded from the infants perspective (2020).

[52] Zhuang, C., She, T., Andonian, A., Mark, M. S. & Yamins, D. Unsupervised learning from video with deep neural embeddings. arXiv preprint arXiv:1905.11954 (2019).

[53] Wong, R. O. Retinal waves and visual system development. Annual review of neuroscience 22, 29–47 (1999).

[54] Zhuang, C., Ding, X., Murli, D. & Yamins, D. Local label propagation for large-scale semi-supervised learning. arXiv preprint arXiv:1905.11581 (2019).

[55] Tarvainen, A. & Valpola, H. Mean teachers are better role models: Weight-averaged consistency targets improve semi-supervised deep learning results. In Advances in neural information processing systems, 1195–1204 (2017).

[56] Wu, M., Zhuang, C., Mosse, M., Yamins, D. & Goodman, N. On mutual information in contrastive learning for visual representations. arXiv preprint arXiv:2005.13149 (2020).

[57] Kar, K., Kubilius, J., Schmidt, K., Issa, E. B. & DiCarlo, J. J. Evidence that recurrent circuits are critical to the ventral streams execution of core object recognition behavior. Nature neuroscience 22, 974–983 (2019).

[58] Nayebi, A. et al. Task-driven convolutional recurrent models of the visual system. In Advances in Neural Information Processing Systems, 5290–5301 (2018).

[59] Bengio, Y., Lee, D.-H., Bornschein, J., Mesnard, T. & Lin, Z. Towards biologically plausible deep learning. arXiv preprint arXiv:1502.04156 (2015).

[60] Zylberberg, J., Murphy, J. T. & DeWeese, M. R. A sparse coding model with synaptically local plasticity and spiking neurons can account for the diverse shapes of v1 simple cell receptive fields. PLoS computational biology 7 (2011).

[61] Kunin, D. et al. Two routes to scalable credit assignment without weight symmetry. arXiv preprint arXiv:2003.01513 (2020).

[62] Mayer, D. L. & Dobson, V. Visual acuity development in infants and young children, as assessed by operant preferential looking. Vision research 22, 1141–1151 (1982).

[63] Gogate, L. J., Bolzani, L. H. & Betancourt, E. A. Attention to maternal multimodal naming by 6-to 8-month-old infants and learning of word–object relations. Infancy 9, 259–288 (2006).

[64] Kachergis, G., Yu, C. & Shiffrin, R. M. Actively learning object names across ambiguous situations. Topics in Cognitive Science 5, 200–213 (2013).

[65] Xu, F. Towards a rational constructivist theory of cognitive development. Psychological review 126, 841 (2019).

[66] Li, M., Liu, F., Jiang, H., Lee, T. S. & Tang, S. Long-term two-photon imaging in awake macaque monkey. Neuron 93, 1049–1057 (2017).

[67] Trautmann, E. M. et al. Accurate estimation of neural population dynamics without spike sorting. Neuron 103, 292–308 (2019).

## Methods References

[68] Lotter, W., Kreiman, G. & Cox, D. Deep predictive coding networks for video prediction and unsupervised learning. arXiv preprint arXiv:1605.08104 (2016).

[69] Zhang, Y. et al. Physically-based rendering for indoor scene understanding using convolutional neural networks. In 2017 CVPR, 5057–5065 (IEEE, 2017).

[70] Olah, C., Mordvintsev, A. & Schubert, L. Feature visualization. Distill (2017). Https://distill.pub/2017/feature-visualization.

